# Jasmonic Acid Oxidases (JAO) define a new branch in jasmonate metabolism towards 11OH-jasmonic acid and its glucosylated derivative

**DOI:** 10.1101/2025.10.09.681447

**Authors:** Thierry Heitz, Valentin Marquis, Julie Zumsteg, Dina Dannawi, Dorian Schutz, Laurence Miesch

**Affiliations:** Institut de Biologie Moléculaire des Plantes (IBMP) du CNRS, Université de Strasbourg, Strasbourg, France; Equipe Synthèse Organique et Phytochimie, Institut de Chimie du CNRS UMR 7177, Université de Strasbourg, Strasbourg, France

**Keywords:** hormone, OH-JA, catabolism, plant stress, 2OGD, Botrytis, wounding

## Abstract

Jasmonates (JAs) occur in plants as a group of related compounds undergoing complex metabolic conversions that shape signatures unique to a given organ or physiological status. Previous studies have shown that several JAs result from direct or indirect catabolism of the master hormonal regulator jasmonoyl-isoleucine (JA-Ile) that controls most jasmonate responses triggered by developmental or environmental cues. Hydroxylation of the precursor jasmonic acid (JA) by Jasmonic Acid Oxidases (JAO) holds a peculiar regulatory function, by attenuating basal JA-Ile formation and action, and promoting growth. Here we reinvestigated biochemically and genetically the nature and origins of hydroxy-JAs and their derivatives in Arabidopsis. Using refined analytical methods and pathway-impaired mutants, we show that JAO enzymes produce exclusively 11OH-JA and are preferentially recruited in response to fungal infection where 11-*O*-Glc-JA accumulates as its main glucosylated derivative. In contrast, mechanical wounding triggers the dominant formation of 12-OH-JA (and its derivatives 12-*O*-Glc-JA and 12-HSO_4_-JA) as a cleavage product of the JA-Ile catabolite 12OH-JA-Ile by the amido hydrolases IAR3 and ILL6. Our results identify the elusive 11OH-JA biosynthetic pathway and provide a revisited picture of JA metabolism where two separate enzyme systems lead to stress-modulated formation of hydroxy-JAs position isomers to attenuate signaling.

## Introduction

Hydroxylated jasmonates (OH-JAs) have been reported as commonly occurring and abundant jasmonic acid (JA) derivatives in many plant species (Miersch et al., 2008). They accumulate under developmental cues in flower organs or in seeds and their synthesis is stimulated in vegetative tissue by environmental stress, typically herbivory or microbial infections (Miersch et al., 2008; Aubert et al., 2015). HO-JA is also readily modified by ‘decorating’ enzymes, such as the sulfotransferase ST2a generating the sulfo-derivative HSO_4_-JA (Gidda et al., 2003), a reaction that is recruited for the modulation of JA signaling and growth during shade avoidance (Fernandez-Milmanda et al., 2020). Alternatively, OH-JA can be readily glucosylated to Glc-*O*-JA by glucosyltransferases of the UGT76 family (Haroth et al., 2019). Although 12OH-JA is an inactive jasmonate in terms of signaling through the canonical JAZ-COI1 co-receptors (Chini et al., 2007; Thines et al., 2007), in pioneering studies, it was described for its tuber-inducing properties in potato (Yoshihara et al., 1989), hence its name tuberonic acid (TA), alongside with its glucoside TAG. In addition, (-)-LCF, a specific enantiomer of TAG, is stimulating motor cells and leaflet closure in the rain tree *Samanea saman* (Nakamura et al., 2011). For many years after its discovery, the biosynthetic pathway of 12OH-JA, the most decribed hydroxy-JA, remained elusive. The first elucidated biosynthetic route derives from two intertwined JA-Ile catabolic pathway, where cytochrome P450 CYP94-generated 12OH-JA-Ile is deconjugated by amido-hydrolases IAR3 and ILL6 to release 12OH-JA (Heitz et al., 2012; Widemann et al., 2013). The second pathway consists in the direct hydroxylation of JA by a small family of 2-oxoglutarate dependent oxidases named Jasmonic Acid Oxidases (JAO) or Jasmonate-induced OXidases (JOX) (Caarls et al., 2017; Smirnova et al., 2017). This metabolic step is bearing a regulatory function as evidenced in Arabidopsis (Smirnova et al., 2017), *Nicotiana attenuata* (Tang et al., 2020) and rice (Ndecky et al., 2025) by repressing JA responses due to JA hydroxylation at the expense of JA-Ile formation. In these successive studies, the product of JAO activity was characterized by standard liquid chromatography coupled to mass spectrometry (LC-MS/MS) and assumed being hydroxylated at the terminal C12 or μ-carbon.

Of importance, early studies in JA metabolism - using mainly gas chromatography coupled to mass spectrometry (GC-MS) analytic methods - have detected the occurrence of 11OH-JA (Helder et al., 1993; Miersch et al., 2008) and its possible derivative 11-Glc-*O*-JA in potato (Matsuura et al., 2001) and *Eschscholtzia californica* cell cultures (Xia and Zenk, 1993), but their biosynthetic origins were not investigated. The elucidation of 11/12OH-JA biochemistry was hampered by the lack of dedicated tools, including resolutive chromatographic conditions, characterized pure reference compounds and genetic resources to validate their *in vitro* formation and *in planta* occurrence. Here we developed chromatographic methods and used characterized synthetic references to discriminate 11OH-JA and 12OH-JA and their derivatives in biological samples by LC-MS/MS analysis and investigate their stress- and pathway-specific formation in challenged Arabidopsis leaves. We show that contrary to previous reports, JAO enzymes produce exclusively 11OH-JA and direct a new branch in JA metabolism. Using mutants impaired either in the deconjugation or in the direct hydroxylation pathway, we found that the 11OH-JA pathway is particularly recruited upon *Botrytis* infection, and that leaf wounding stimulates the preferential formation of 12OH-JA and its glycosylated and sulfated derivatives. Our results provide an updated view of OH-JA diversity and illustrate a new complexity in signal attenuation architecture within the jasmonate metabolic pathway.

## RESULTS

### Impairing hydroxy-JA biosynthetic pathways modifies distinctly JAs homeostasis in leaf stress specific patterns

To study the accumulation of OH-JA and its relationship to global jasmonate signatures, we submitted WT Arabidopsis plants to mechanical wounding - a proxy for herbivory - or to infection with the necrotrophic fungus *Botrytis cinerea* (Bc), two biotic assaults that powerfully trigger jasmonate metabolism and responses (Aubert et al., 2015; Marquis et al., 2020) and established quanTtaTve JAs profiles. In non stimulated leaves, basal OH-JA levels were close to detection limit; in contrast, its accumulation was strongly stimulated 3.5 h post wounding or three days after fungal inoculation (Fig. 1A upper panel). In parallel to WT, we setup the analysis of three genetic backgrounds impaired in either one or both known HO-JA biosynthetic pathways : *2ah*, a double mutant deficient for amidohydrolases (AH) IAR3 and ILL6 cleaving JA-amino acid conjugates (Widemann et al., 2013; Heitz et al., 2019; Heitz, 2020; Marquis et al., 2020); *3jao*, a triple mutant impaired in *JAO2, JAO3, and JAO4* genes encoding JA oxidase activity in leaves (Marquis et al., 2022); and *5ko*, a newly introduced quintuple mutant line combining both defects. The mutations had distinct impacts on jasmonate profiles with regards to the nature of the stress. Upon wounding, JA was strongly overaccumulated in JAO-deficient lines (Supplemental Figure S1A). In contrast, HO-JA level was reduced in *2ah* and more significantly in *5ko*, and this trend mirrored an overaccumulation of its precursor 12OH-JA-Ile and downstream product 12COOH-JA-Ile in these genotypes. To try to understand why wounded *5ko* leaves still contained substantial amounts of OH-JA despite of having both pathways impaired for their major biosynthetic genes, we hypothetized that related genes that were not mutated in these plant lines may have a compensatory elevated expression in the mutants under study: when we monitored the expression of *JAO1 –* a nearly non-expressed gene in leaves (Smirnova et al., 2017) *-* and members of the amido-hydrolase family including *ILL1, ILL2, ILL3, ILL5* and *ILR1*, only *ILL5* transcripts were overaccumulated in *2ah* and *5ko*. ILL5 is a close relative to IAR3 (LeClere et al., 2002) but it is a pseudogene in Col0 with a 7 nt deletion in coding sequence (Widemann et al., 2013), therefore it cannot account for 12OH-JA production via 12OH-JA-Ile cleavage.

**Figure 1.**
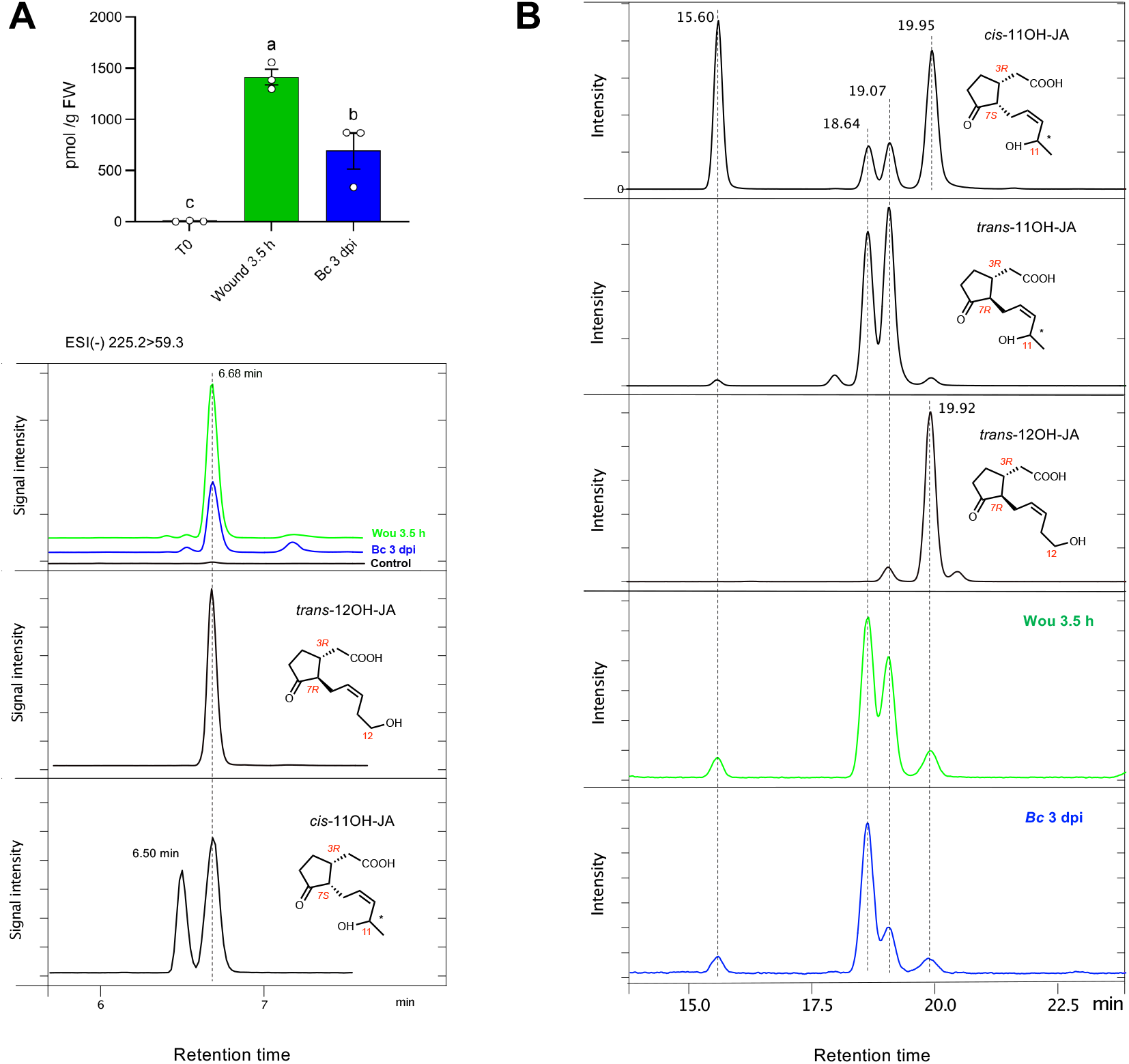
Highly resolutive chromatographic conditions separate OH-JA accumulated in response to wounding or infection into distinct isomers. (A) Upper panel: quantification of OH-JA in untreated, wounded (3.5 h) or infected leaves (3 days post-inoculation, dpi) of 6 week-old plants. JAs were extracted, analyzed and quantified by standard LC-MS/MS (see Methods). Histograms are means ± SEM of 3 biological replicates. Statistical significance was assessed by one-way ANOVA followed by Tukey post-hoc test with a 95% confidence interval. Different letters above boxplots indicate genotypes that are significantly different (P< 0.05). Lower panels : chromatographic traces of OH-JA (ESI-) 225.2>59.3 in standard methanol gradient with 100 mm column. Traces show co-elution of signal with references of *trans*-12OH-JA and one peak of *cis*-11OH-JA at 6.68 min. (B) Chromatographic analysis of OH-JAs with 150 mm column and 9-11% acetonitrile gradient. Preparations of *cis*-11OH-JA, *trans*-11OH-JA and *trans*-12OH-JA were analyzed along with extracts from wounded (wou 3.5 h) or infected (Bc 3 dpi) leaves.

In infected leaves, all Ile-conjugates were more abundant in *2ah* (Supplemental Fig. S1B), consistent with a previous report (Marquis et al., 2020), and OH-JA was strongly diminished by 77% in both JAO-impaired lines. When antifungal resistance was assessed, *2ah* displayed WT-size fungal lesions as shown previously (Marquis et al., 2020), but *3jao* was more resistant (Supplemental Fig. S1C), corroborating a phenotype described for a double *jao2 jao4* line (Smirnova et al., 2017) and a quadruple *jao/joxQ* mutant (Caarls et al., 2017). Furthermore, *5ko* plants phenocopied *3jao* in this assay, indicating the prevalent impact of the JAO pathway on resistance to *Botrytis*. Collectively, these results illustrate that hydroxy-JA biosynthetic pathways are important drivers of the global dynamics of JAs upon leaf stress in Arabidopsis.

### Wounding and *Botrytis* infection stimulate differential accumulation of 11- and 12-hydroxy-jasmonic acid in Arabidopsis leaves

In our standard liquid chromatography conditions using a methanol gradient, HO-JA from stimulated leaf extracts co-eluted at 6.68 min with a synthetic standard of (*3R, 7S*)-12OH-JA (also referred to as *trans*-12OH-JA). To explore the potential accumulation of 11OH-JA in plants, we utilized a reference obtained by chemical synthesis of *3R,7S*-11OH-JA (also referred to as *cis*-11OH-JA) as described in the accompanying paper (Matsumoto et al., 2025). The synthetic sample was resolved as two separated peaks at 6.50 and 6.68 min, the latter co-eluting with the *trans-*12OH-JA reference (Fig. 1A lower panel). This profile corresponds to the separation of two 11OH-JA stereoisomers at C11. We next developed a more resolutive chromatographic method, using a longer column and a mild acetonitrile gradient. Under these conditions, the sample of synthetic 11OH-JA separated into two main peaks eluting at 15.60 and 19.95 min, and two minor peaks at 18.64 and 19.07 min (Fig. 1B upper panel). This profile indicates that the *cis*-11OH-JA forms (15.60 min and 19.95 min) have partially isomerized to the more stable *trans*-11OH-JA forms (18.64 min and 19.07 min) during long-distance transport or analysis and therefore appeared as a mixture of four stereoisomers at carbon 7 and 11 (C7 and C11). This was confirmed by analysis of a genuine sample of synthetic *trans*-11OH-JA that was highly enriched in the central peaks eluting at 18.64 min and 19.07 min (Fig. 1B second panel). *trans*-12OH-JA reference eluted as a major peak at 19.92 min and a minor peak at 19.07 min was assigned the less stable *cis*-12OH-JA (Fig. 1B third panel). In extracts of wounded or infected leaves (Fig. 1B fourth and fifth panels), four peaks were also detected, but here, the two central peaks were dominant. We assumed that in plant samples, compounds had mostly isomerized during workup and storage from the enzyme-generated *cis*-isomers to the more stable *trans*-isomers. These results define new analytical conditions to separate *trans*-hydroxy-JAs that differ at the C11 or C12 hydroxylation position and allow to examine their occurrence in Arabidopsis stress-responding leaves. The profiles suggest that both stresses trigger the accumulation of 11OH- and 12OH-JA in leaves, but 11OH-JA appears as the predominant isomer in response to *Botrytis* infection, raising the question of.

### Jasmonic Acid Oxidases produce predominantly 11OH-JA

HO-JA was previously reported to be formed by two separate biosynthetic pathways, either by the CYP94-AH pathway acting on 12OH-JA-amino-acid conjugates (Widemann et al., 2013; Widemann et al., 2015), or by direct JA hydroxylation catalyzed by JAO activity (Caarls et al., 2017; Smirnova et al., 2017). Because the JAO enzymatic product was so far assumed to be the commonly occurring 12OH-JA, on the basis of routine separation methods but without formal identification, we re-investigated the JA hydroxylation product generated by recombinant JAO enzymes using the newly developed chromatographic tools. As shown in Figure 2, the resolutive conditions revealed that recombinant Arabidopsis JAO2, 3, 4 and the recently characterized OsJAO2 and OsJAO3 from rice (Ndecky et al., 2025) produce near exclusively 11OH-JA (Fig. 2). The presence of two peaks coeluting with the *trans*-11OH-JA reference suggests that JAO enzymes generate both *R* and *S* stereoisomers at C11.

**Figure 2.**
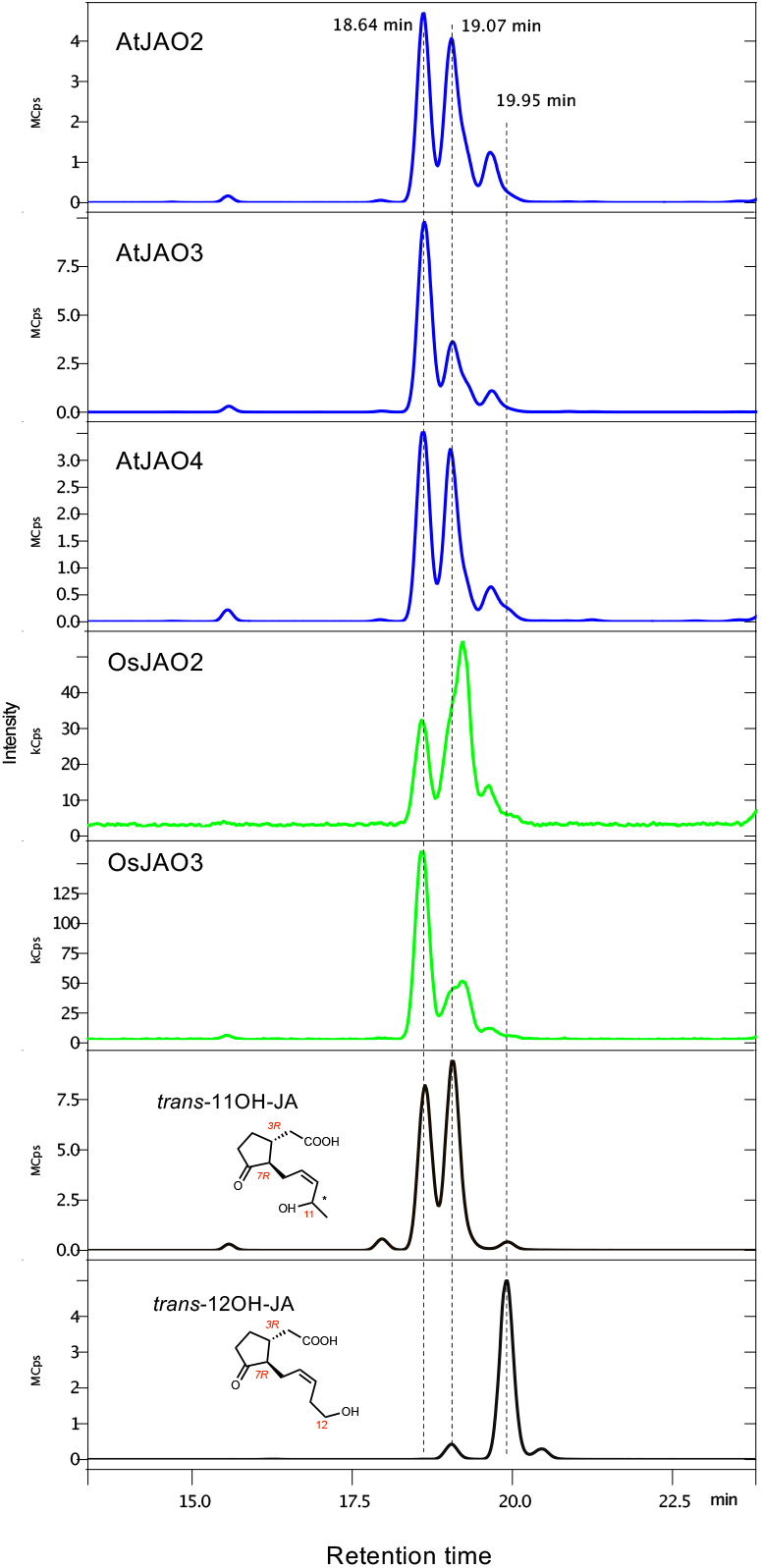
JAO proteins from Arabidopsis and rice produce 11OH-JA. Indicated his-tagged JAO proteins were expressed in bacteria, affinity-purified and incubated with 50 µM JA in presence of the co-substrate 2-oxoglutarate for 40 min at 30°C. Reaction mixture was analyzed by LC-MS/MS using a 150 mm column and a 9-11% acetonitrile gradient. Traces show analysis of Arabidopsis enzymes AtJAO2, AtJAO3, AtJAO4 and rice enzymes OsJAO2 and OsJAO3 along with synthetic references 11OH-JA and 12OH-JA.

### Deconjugation and direct oxidation pathways produce different position OH-JA isomers with stress-specific signatures

The two biochemical pathways producing OH-JA are both transcriptionally upregulated by biotic stress in leaves (Kitaoka et al., 2011; Koo et al., 2011; Heitz et al., 2012; Widemann et al., 2013; Aubert et al., 2015; Smirnova et al., 2017). The preferential recruitement of the deconjugation pathway (CYP94-AH) upon wounding and of the direct hydroxylation (JAO) pathway by fungal infection respectively was suggested previously based on global JAs profiling (Aubert et al., 2015; Smirnova et al., 2017). This distinctive feature provides the opportunity to assess genetically the contributions of each hydroxylation route in response to these two types of assaults. To this end, we compared the accumulation of HO-JAs in wounded or infected leaves of WT, *2ah, 3jao* and *5ko* genetic backgrounds. Strikingly, each of the four genotype presented a specific pattern of OH-JA accumulation upon wounding (Fig. 3A left), with respect to the three main peaks at 18.64, 19.07 and 19.95 min. Major peaks in WT (18.64 and 19.07 min) matched those of the *trans*-11OH-JA reference and the major peak (19.95 min) of the 12OH-JA reference. All peak areas were quantified in biological replicates and their statistical differences were assessed (Fig. 3A right panels). In *2ah*, abundance of peak 1 (18.64 min), corresponding to an 11OH-JA isomer, was not affected relative to WT but peak 2 (19.07) was diminished by half in *2ah* and in *5ko*. In contrast, peak 1 was lost in *3jao* and in *5ko*. Finally, peak 3 (19.95 min) that co-elutes with *trans*-12OH-JA reference was negatively and positively affected by *2ah* and *3jao* mutations respectively. These patterns support a scenario where 11OH-JA biosynthesis is JAO-dependent and AH activity generates 12OH-JA. They imply that the peak at 19.07 min in plant extracts may be a mixture of 11OH- and 12OH-isomers. Its increased abundance in *3jao* suggests an overstimulation of 12OH-JA synthesis due to enhanced JA-Ile signaling leading to upregulation of IAR3/ILL6 and AH activity as reported previously (Smirnova et al., 2017; Marquis et al., 2022). Therefore, despite of co-eluting with a peak of *trans*-11OH-JA reference, peak 2 of plant extract probably contains the *cis-* isomer of 12OH-JA.

**Figure 3.**
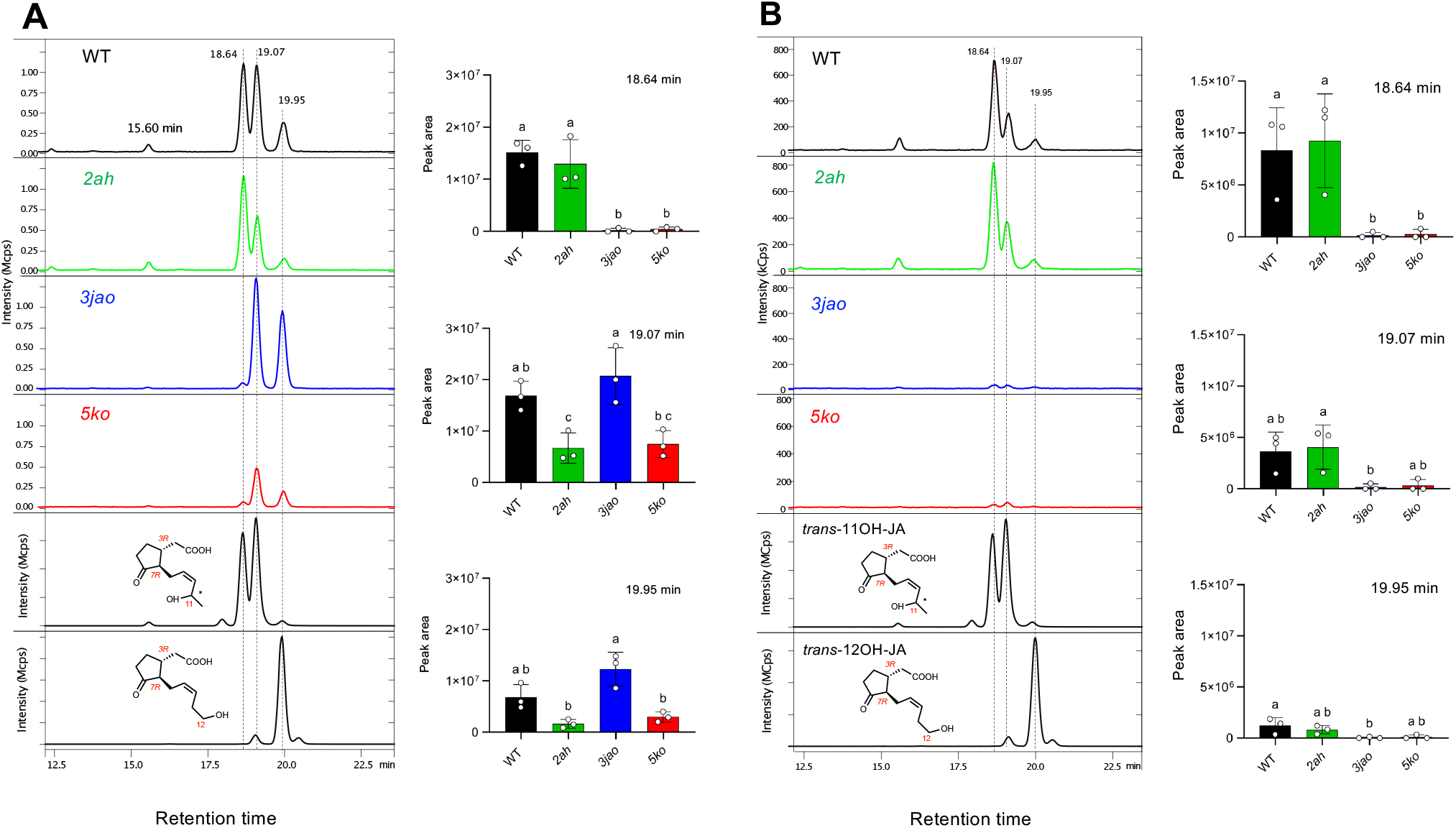
High-resolution LC-MS/MS analysis of OH-JAs in WT plants and mutant lines impaired in the deconjugation (*2ah*), or the direct hydroxylation (*3jao*) or both (*5ko*) pathways upon (A) leaf wounding or (B) fungal infection. Left panels in (A) and (B): high resolution representative chromatograms of plant extracts obtained with a 9-11% acetonitrile gradient along with *trans*-11OH-JA and 12OH-JA synthetic references. Histograms show means ± SEM of 3 biological replicates indicated with circles. Statistical significance was assessed by one-way ANOVA followed by Tukey post-hoc test with a 95% confidence interval. Different letters above boxplots indicate genotypes that are significantly different (P< 0.05). Two independent experiments were performed with similar results.

A similar analysis was conducted with extracts from fungus-infected leaves. Here, 11OH-JA (peak 1) was found to largely dominate in WT (Fig. 3B left panel), consistent with our previous report of preferential recruitment of the JAO pathway in response to *Botrytis* (Smirnova et al., 2017). This profile remained unchanged in *2ah*, illustrating the marginal contribution of AH activity to the accumulation of OH-JA upon necrotrophic infection. Most strikingly, infected *3jao* and *5ko* leaves were essentially depleted of both position isomers of OH-JA. These results indicate that two separate JA hydroxylation pathways co-exist in Arabidopsis : the CYP-AH pathway that generates 12OH-JA and the JAO pathway that produces 11OH-JA are both upregulated upon wounding, but infection recruits mostly the JAO pathway.

### JAO activity provides the precursor for biosynthesis of 11-*O*-Glc-JA

12OH-JA commonly occurs as abundant glucosylated and sulfated forms in most plant species analyzed (Miersch et al., 2008), with decorations introduced on the ω-hydroxy function by UGT76E1 and ST2a enzymes respectively (Gidda et al., 2003; Haroth et al., 2019). In parallel, despite of the long-standing identification of 11-*O*-Glc-JA in suspension cultures of *E. californica* (Xia and Zenk, 1993) or in potato leaflets (Matsuura et al., 2001), the biosynthetic route for this derivative is not elucidated. To explore the mode of formation of 11OH-JA derivatives and distinguish them from the steps proceeding through 12OH-JA, we first synthetized enzymatically glucosylated derivatives from synthetic 11OH-JA and 12OH-JA substrates using recombinant UGT76E1 from Arabidopsis expressed in bacteria. Both expected compounds were obtained, but UGT76E1 glucosyl-transferase glycosylated 11OH-JA about 7 fold less efficiently than 12OH-JA (Fig. 4A left fifth and sixth panels), in agreement with a previous report (Haroth et al., 2019). 11-*O*-Glc-JA (6.05 min) and 12-*O*-Glc-JA (6.26 min) reference compounds were sufficiently separated in standard MeOH-gradient chromatography to examine their occurrence in plant extracts. In wounded leaves, 12-*O*-Glc-JA largely dominated the WT profile (Fig. 4A), and its abundance was strongly decreased in *2ah* and moderately in *5ko*, while being unaffected in *3jao*. Conversely, 11-*O*-Glc-JA (6.05 min) and a second minor peak at 6.66 min were significantly reduced in *JAO*-deficient lines. These results indicate that upon wounding, 12-*O*-Glc-JA is the predominant glucosylated form accumulated, and its biosynthesis is dependent on 12OH-JA generation by the CYP94-AH pathway.

**Figure 4.**
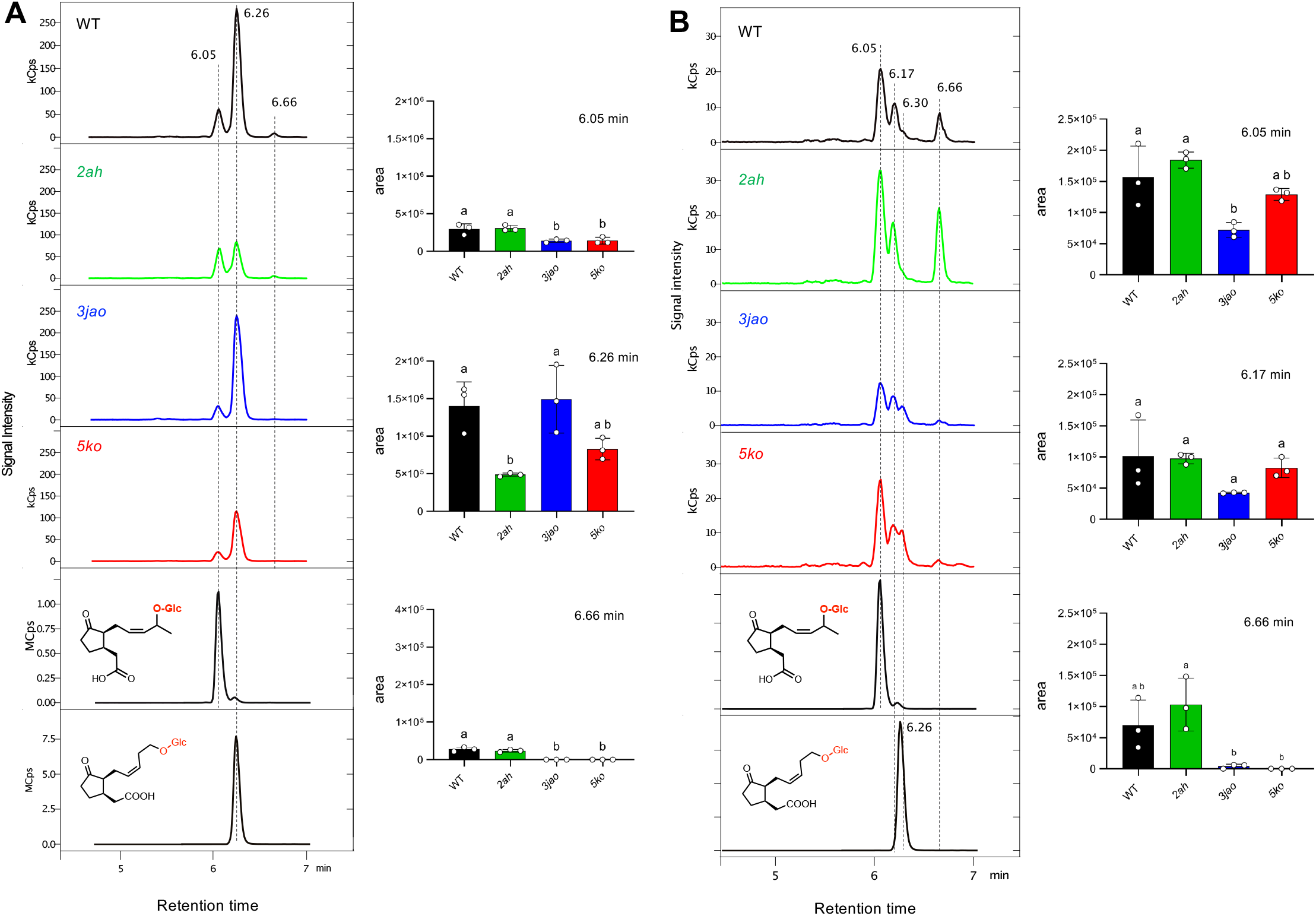
LC-MS/MS analysis of Glc-*O*-JAs in WT plants and mutant lines impaired in the deconjugation (*2ah*), or the direct hydroxylation (*3jao*) or both (*5ko*) pathways upon (A) leaf wounding or (B) fungal infection. Left panels in (A) and (B): representative chromatograms of plant extracts analyzed by standard LC-MS/MS using a methanol gradient along with enzymatically generated Glc-*O*-JA references. Histrograms show means ± SEM of 3 biological replicates indicated with circles. Statistical significance was assessed by one-way ANOVA followed by Tukey post-hoc test with a 95% confidence interval. Different letters above boxplots indicate genotypes that are significantly different (P< 0.05). Two independent experiments were performed with similar results.

A mirrored situation was evidenced in fungus-infected leaves. Here, 11-*O*-Glc-JA (6.05 min) was dominating the profile in WT and *2ah* extracts, and its abundance was reduced by half in *3jao* (Fig. 4B). The unidentified 6.66 min compound was here more pronounced than upon wounding, and disappeared in lines bearing the *3jao* mutations. This signal may be tentatively assigned to *cis*-11-Glc-*O*-JA. Of note, a compound eluted slightly earlier than 12-Glc-*O*-JA reference at 6.17 min in extracts from WT leaves and its abundance was barely affected in *2ah*. In addition, an extra peak at 6.30 min was specific to *3jao* and *5ko* genotypes. These data show that in response to fungal infection, JAO-dependent synthesis of 11-*O*-Glc-JA dominates and the depletion of JAO activity results in additional peaks. Surprisingly, substantial amounts of Glc-*O*-JA subsisted in *5ko* that split into a complex chromatographic pattern shared with *3jao*.

We next examined the formation of HSO_4_-JA, the sulfated derivative of OH-JA. HSO_4_-JA signal was about 2000 fold less intense when recombinant ST2a sulfotransferase was incubated with 11OH-JA compared to 12OH-JA (Fig. 5A left panel), confirming an initial report that 11OH-JA is a poor substrate for ST2a (Gidda et al., 2003). The two enzymatic products where essentially separated, eluting at 6.02 min (11-isomer) and 6.16 min (12-isomer). When HSO_4_-JA was analyzed in plants with respect to genotypes and stimulus, only the later-eluting peak was recorded, suggesting that only 12-HSO_4_-JA was formed *in planta* under our conditions. In wounded leaves, opposite trends were recorded in mutant backgrounds, with an enhanced abundance of 12-HSO_4_-JA in *3jao*, a decrease in *2ah*, and near WT levels in *5ko*. This pattern is consistent with the CYP94-AH pathway feeding 12OH-JA substrate towards 12-HSO_4_-JA formation by ST2a in wounded leaves. Surprisingly, no significant impact of the mutations was evidenced for this derivative upon *Botrytis* infection, leaving open the possibility of alternative route(s) for its formation in this material.

**Figure 5.**
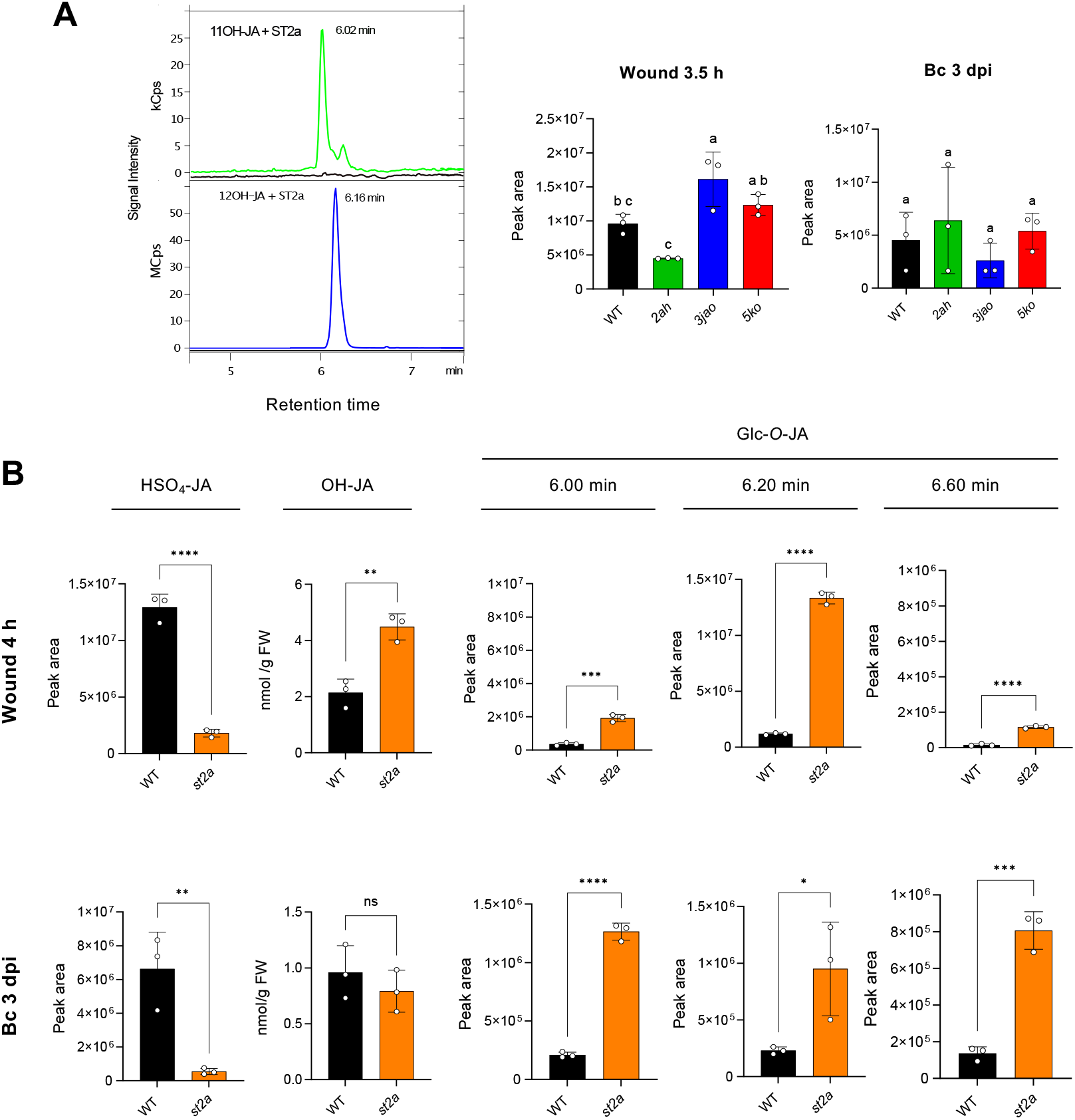
Interaction of OH-JA glucosylation and sulfation pathways. (A) Left panel : LC-MS/MS analysis of HSO_4_-JA obtained after incubation of 11OH-JA or 12OH-JA with ST2a enzyme. Right panels : accumulation of HSO4-JA upon wounding (3.5 h) of fungal infection (3 dpi) in WT and mutant lines impaired in the deconjugation (*2ah*), or the direct hydroxylation (*3jao*) or both (*5ko*) pathways. Histograms show means ± SEM of 3 biological replicates indicated with circles. Statistical significance was assessed by one-way ANOVA followed by Tukey post-hoc test with a 95% confidence interval. Different letters above boxplots indicate genotypes that are significantly different (P< 0.05). Two independent experiments were performed with similar results. (B) Quantification of HSO_4_-JA, OH-JA and Glc-*O*-JA upon wounding or fungal infection (Bc) in WT and *st2a* plants. Histograms show means ± SEM of 3 biological replicates indicated with circles. Statistical significance was assessed in (A) by one-way ANOVA followed by Tukey post-hoc test. Different letters above boxplots indicate genotypes that are significantly different (P< 0.05). In (B),

Finally, we investigated the interaction between sulfation and glucosylation pathways. As expected, 12-HSO_4_-JA levels were strongly decreased in a *st2a* mutant after either wounding or infection challenge (Fig. 5B top left panels). Total OH-JA levels were doubled in wounded *st2a*, probably because 12OH-JA was no longer consumed by ST2a activity. In contrast, OH-JA levels were not altered in infected *st2a*, further suggesting that ST2a does not turnover significantly the 11OH-JA isomer dominating in this material. However in *st2a*, levels of three isomers of Glc-*O*-JA were enhanced by both stimuli relative to WT. Interestingly, the strongest overaccumulation was observed for 12-*O*-Glc-JA after wounding, and for 11-*O*-Glc-JA upon infection, likely reflecting the stress-specific redirection of metabolic flux.

## DISCUSSION

Hydroxy-jasmonic acid (OH-JA) has been identified as a commonly occurring JA oxidation product along with its glucoslated and sulfated derivatives as part of larger panels of plant regulatory metabolites (Gasperini and Howe, 2024). OH-JA, TAG and 12-HSO_4_-JA can build-up as abundant stored pools (Miersch et al., 2008); in several cases, they have been associated with biological responses including growth under shade or warm temperatures, pathogen defense or nyctinastic leaf-folding (Nakamura et al., 2011; Patkar et al., 2015; Ueda et al., 2015; Fernandez-Milmanda et al., 2020; Zhu et al., 2021), or their formation represents a regulatory sink that modulates JA-Ile formation and signaling (Smirnova et al., 2017; Marquis et al., 2022; Ndecky et al., 2025). Since the early reports on the existence of 11OH-JA isomers and glycosylated forms several decades ago (Xia and Zenk, 1993; Matsuura et al., 2001), the mode of formation of these latter species and how they compare to 12OH-JA forms has remained undefined. Despite of its growing biological relevance to plant adaptation to environmental constraints, the elucidation of OH-JA isomer biochemistry was hampered by the lack of some characterized references, adequate separation methods, and genetic tools to track their *in planta* formation.

Here we have assembled a toolbox to fulfill these requirements and re-evaluated the occurrence of 11OH- and 12OH-JA and derivatives in stressed Arabidopsis leaves, with particular focus on the contribution of JAO enzymes. While the 12OH-JA reference is at hands in many laboratories and commercially available, a proper 11OH-JA sample needed to be re-synthetized chemically and characterized. Owing to the full synthesis and characterization of 11OH-JA stereoisomers by Matsumoto et al., (2025), combined to our methodological developments in chromatography allowed to separate and identify major isomers in biological samples. Driven by our early suspicion that AtJAO2 product was structurally distinct from the 12OH-JA reference, we established that JAO/JOX proteins from Arabidopsis and from rice direct the formation of 11OH-JA and its derivatives rather than 12OH-JA as commonly accepted. The existence of this novel branch in JA-metabolism was supported by *in vitro* enzymology data and the analysis of plants submitted leaf stress. Products of recombinant Arabidopsis AtJAO2, AtJAO3 and AtJAO4 and the recently characterized OsJAO2 and OsJAO3 from rice (Ndecky et al., 2025) matched the *trans-*11OH-JA rather than the *trans*-12OH-JA reference. This result sheds new light on an overlooked aspect in initial characterization of JAO/JOX enzymes (Caarls et al., 2017; Smirnova et al., 2017; Tang et al., 2020). We next took advantage of the preferential stimulation of the CYP94-AH and JAO hydroxylation pathways by leaf wounding and fungal infection respectively, two stresses that strongly stimulate JAs metabolism in leaves, but these different physiological contexts and hormonal interactions generate unique JAs signatures and signaling outputs (Aubert et al., 2015; Widemann et al., 2016). For example, the finding that Botrytis induces the F-box protein BFP1 directing the proteolytic degradation of JAO2 (Zhang et al., 2024) illustrates the importance of JA hydroxylation in the plant-pathogen arms race. The use of mutant lines impaired in either one or both hydroxylation pathways illustrated that *2ah, 3jao* or *5ko* mutations had profound but distinct consequences on steady-state levels of several JAs, reflecting different metabolic fluxes within the JAs biochemical pathway during simulated herbivory and fungal infection.

Most importantly, this set of plant genetic backgrounds revealed clearly the generation of stress-specific blends of OH-JA forms in Arabidopsis leaves. In WT, wounding triggers comparable accumulation of 11OH-and 12OH-JA at 3.5 h after injury, but these proportions may vary at other timepoints. *3jao* or *2ah* mutations diminished or even suppressed 11OH-JA or 12OH-JA specific signal, assigning unambiguously the synthesis of 11-OH-JA to JAO activity and the origin of 12OH-JA to deconjugation of CYP94-generated 12OH-JA-Ile by AH activity. Surprisingly, in wounded *5ko* leaves, residual 12OH-JA was observed. Given that besides IAR3 and ILL6 no AH is known to cleave 12OH-JA-aminoacid conjugates based on *in vitro* assays (Widemann et al., 2013; Zhang et al., 2016), the origin of this pool of 12OH-JA remains unknown, and this observation deserves further investigation. Conversely, in *Bc*-infected leaves, the accumulation of JAO-dependent 11OH-JA was largely predominant, and curiously, minor 12OH-JA was also suppressed by JAO deficiency. Additional levels of interactions between the two hydroxylation pathways may result in currently unexplained outcomes.

Because they represent important and abundant end-products that may drive biological responses mediated by OH-JAs (Fernandez-Milmanda et al., 2020; Zhu et al., 2021), we investigated genetically the formation of the derivatives Glc-*O*-JA and HSO_4_-JA, taking advantage of enzymatically generated references from synthetic OH-JAs. As expected from the newly established OH-JA formation pathways, 12-Glc-*O*-JA (TAG) was predominantly formed upon wounding via AH-mediated deconjugation of 12OH-JA-Ile, with no contribution of the JAO pathway. Again, the *2ah* mutation did not cancel the accumulation of 12-Glc-*O*-JA. Despite of the absence of reported *in vitro* 12OH-JA-Ile cleaving activity for other members of the AH family (LeClere et al., 2002; Zhang et al., 2016), we cannot exclude that residual AH activity in *2ah* and *5ko* lines releases 12OH-JA for further metabolization. In contrast, upon infection, mostly 11-Glc-*O*-JA was formed from JAO-generated substrate. Finally, we found only evidence for sulfation of 12OH-JA by ST2a, irrespective of the stimulus applied, in accordance with the *in vitro* preference of this enzyme for 12OH-JA substrate. Moreover, impairement of the sulfation in *st2a* pathway redirected metabolic flux towards enhanced formation of different isomers of Glc-*O*-JA, illustrating the competition of glucosylation and sulfation steps for common substrates.

In conclusion, our results provide a updated view of the JA metabolic pathway, by elucidating the elusive formation of 11OH-JA (Fig. 6). At odds with conclusions from previous reports, we demonstrate that JAO enzymes direct the exclusive formation of 11OH-JA in Arabidopsis and likely in rice. These findings call for the use of more resolutive analysis conditions in future studies to allow accurate identification of isomers and harness the proper interpretation of OH-JA profiles. Here, the combination of genetic and fine chromatographic analysis illustrate the recruitment of separate 11OH- and 12OH-JA biosynthetic pathways and distinct glucosylation steps upon fungal infection and wounding in Arabidopsis leaves. In contrast sulfation seems to occur only on the 12OH-JA isomer. These new metabolic routes expand the metabolic flexibility and regulatory potential to adjust finely JA-Ile dynamics and action. Duplication of attenuation mechanisms upstream (JAO) and downstream (CYP94-AH) of the master regulator JA-Ile ensures robustness and safeguards plants from overamplification of the response. Despite of these advances, some of the metabolic profile characteristics recorded here remain unexplained, suggesting the existence of yet unknown conversion circuits or regulatory checkpoints. For example, even though wounding generates the preferential formation of 12OH-JA, the *3jao* mutation and not *2ah*, elevates precursor JA accumulation. Conversely, fungal infection triggers mostly the formation of 11OH-JA, but its impairement in *3jao* does not lead to enhanced JA accumulation. Future investigations of metabolic fluxes should clarify these and other uncertainties as well as their biological implications.

**Figure 6.**
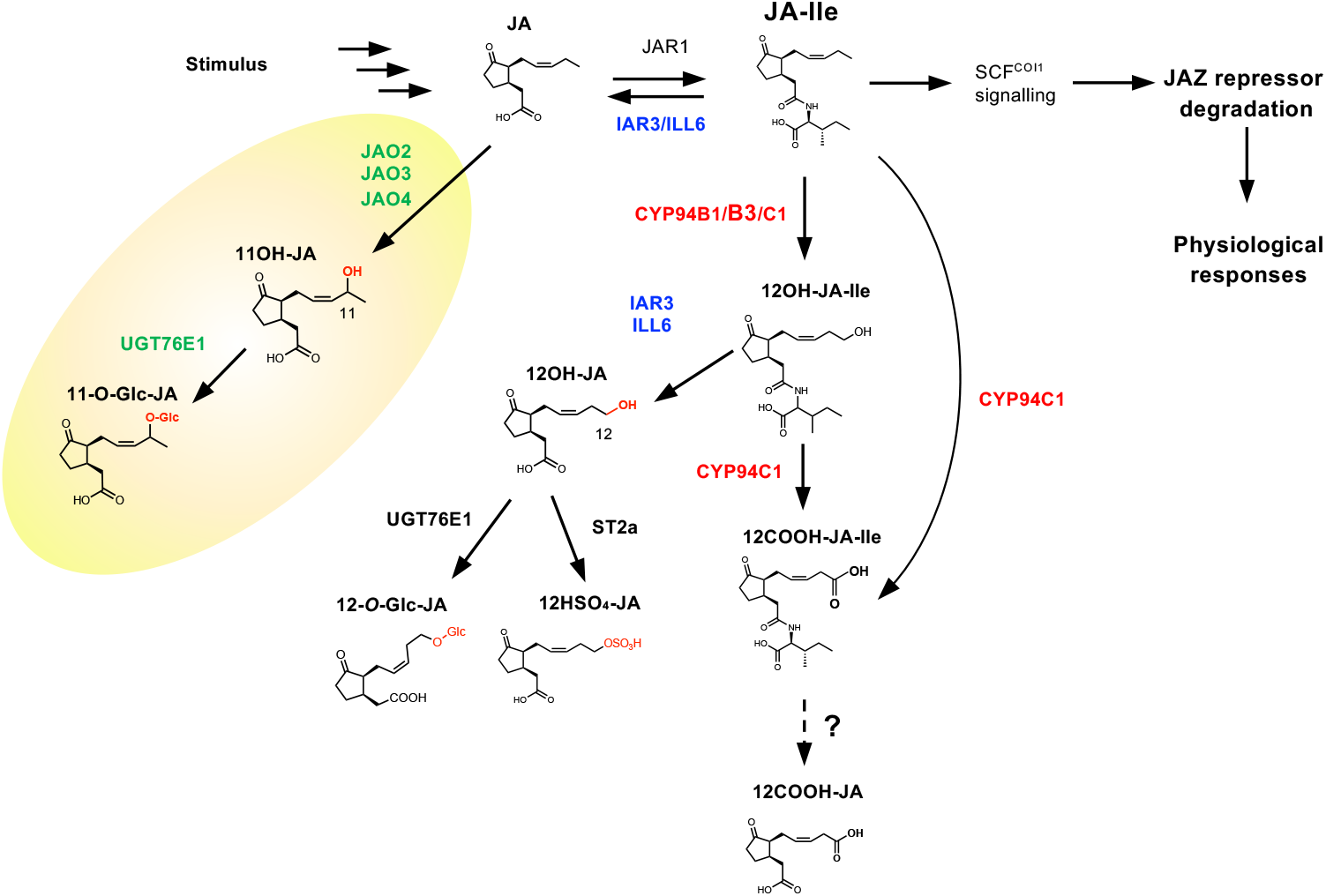
Updated model of OH-JA formation in the general JA metabolic pathway in Arabidopsis leaves. Jasmonic acid is formed upon a stimulus and activated by conjugation into the major conjugate JA-Ile that triggers physiological responses via COI-mediated JAZ repressor degradation. JA-Ile catabolism provides an indirect route to 12OH-JA formation by successive oxidation to 12OH-JA-Ile by cytochromes P450 of the CYP94 family, and cleavage by the amido hydrolases IAR3 and ILL6. This pathway is preferentially activated upon leaf wounding, and 12OH-JA is further modified by glucosylation (UGT76E enzymes) or sulfation (ST2a). Upstream of JA-Ile formation, JA is directly oxidized by JA oxidases (JAO), providing a conserved regulatory node for basal JA-Ile signaling. The JAO pathway directs the formation of the distinct isomer 11OH-JA and its glucosylated derivative, and is particularly recruited upon fungal infection. The renewed metabolic grid illustrates the existence of parallel JA hydroxylation pathways.

## MATERIALS AND METHODS

### Plant growth and treatments

*Arabidopsis thaliana* genotypes used were in the Col-0 ecotype and were grown under a 12 h light/12 h dark photoperiod in a growth chamber. The individual T-DNA insertion lines including the *st2a-2* (GK149G04) line were obtained from the Nottingham Arabidopsis Stock Center (NASC). The *3jao* line (Marquis et al., 2022) was obtained by crossing the alleles *jao2-1* (SALK_206337C), *jao3-1* (SAIL_861E01) and *jao4-2* (SAIL_268_B05). The *2ah* line (Marquis et al., 2020) was obtained by crossing the lines *iar3-5* (SALK_069047) and *ill6-2* (SALK_024894C). The quintuple *5ko* line was obtained by crossing the *3jao* line with a double *iar3-5 ill6-1* (GK412-E11).

*B. cinerea* inoculation and resistance scoring were as described in Aubert et al., (2015). For mechanical wounding experiments, 5 or 6 fully expanded leaves from 6 week-old plants were wounded three times across mid-vein with a hemostat. At increasing time points following damage, leaf samples were quickly harvested and flash-frozen in liquid nitrogen before storing at −80°C until analysis.

### Jasmonate profiling

Jasmonates were separated, identified and quantified by ultra high performance liquid chromatography coupled to tandem mass spectrometry (UHPLC-MS/MS). Frozen plant material (in the 50-60 mg range) was precisely weighed and extracted with 8 volumes of ice-cold extraction solution (MeOH:water:acetic acid 80:19:0.5, v/v/v) containing d5-JA (Olchemim, Olomuc) and 9,10-dihydro-JA-Ile as internal standards for workup recovery. Grinding was performed with a glass-bead Precellys tissue homogenizer (Bertin Instruments, France) in 2 mL screw-capped tubes. After 30 min incubation at 4°C on a rotating wheel, homogenates were cleared by centrifugation before 1.8x concentration of supernatants under a stream of N_2_ and overnight conservation at −20°C. After a second centrifugation step, extracts were submitted to quantitative LC-MS/MS analysis on an EvoQ Elite LC-TQ (Bruker) equiped with an electrospray ionisation source (ESI) and coupled to a Dionex UltiMate 3000 UHPLC system (Thermo). Five µL plant extract were injected. For routine analysis, chromatographic separation was achieved using an Acquity UPLC HSS T3 column (100 x 2.1 mm, 1.8 µm; Waters) and pre-column. The mobile phase consisted of (A) water and (B) methanol, both containing 0.1 % formic acid. The run started by 2 min of 95 % A, then a linear gradient was applied to reach 100 % B at 10 min, followed by isocratic run using B during 3 min. Return to initial conditions was achieved in 1 min, with a total run time of 15 min. For position-specific analysis of 11OH-JA and 12OH-JA, chromatographic separation was achieved using an Acquity UPLC UPLC HSS T3 column (150 x 2.1 mm; 1.8 µm; Waters) and pre-column. Samples were carried through the column following a gradient of solvent A (H20; 0.1% formic acid) and B (acetonitrile; 0.1% formic acid) starting with 9% B for 1 min, reaching 11% B at 41 min with a convex curve 6, returning to 9% B in 1 min and equilibrating for 1 minute afterwards, for a total run time of 43 min. The columns were operated at 35 °C with a flow-rate of 0.30 mL min^−1^.

Nitrogen was used as the drying and nebulizing gas. The nebulizer gas flow was set to 35 L h^−1^, and the desolvation gas flow to 30 L h^−1^. The interface temperature was set to 350 °C and the source temperature to 300 °C. The capillary voltage was set to 3.5 kV, the ionization was in positive or negative mode. Low mass and high mass resolution were 2 for the first mass analyzer and 2 for the second. Data acquisition was performed with the MS Workstation 8 for the mass spectrometry and the liquid chromatography was piloted with Bruker Compass Hystar 4.1 SR1 software. The data analysis was performed with the MS Data Review software. Absolute quantifications were achieved by comparison of sample signal with dose-response curves established with pure compounds and recovery correction based on internal standard signal. The transitions were, in negative mode: JA 209.3>59.3; JA-Ile 322.3>130.2; 11/12OH-JA 225.2>59.3; 12OH-JA-Ile 338.3>130.2; 12COOH-JA-Ile 352.2>130.1; 12COOH-JA 239.2>59.3; Glc-*O*-JA; 387.3>59.3; HSO_4_-JA 304.9>97.1.

### Protein expression and enzyme assays

Total RNA was extracted from plant leaves harvested 3.5 h post-wounding with TRI reagent (Molecular Research Center). One microgram of RNA was reverse transcribed using SuperScript IV reverse transcription system (Thermo Fisher Scientific). cDNAs coding for UGT76E1 (At5g59580) and ST2a (At5g07010) were PCR-amplified using Phusion Taq Polymerase (Thermo Fisher Scientific) and inserted into pDONRzeo by BP recombination using Gateway cloning. Inserts were recombined in pHMGWA expression plasmid (Busso 2005). Sequence-verified plasmids were next transferred in the *Escherichia coli* Rosetta 2 pLys strain. Bacteria were grown at 37°C in LB medium at 250 rpm up to optical density (A_600_) of 0.5. Expression of recombinant proteins was induced by adding IPTG 0.5 mM and further incubated under shaking at 20°C for 5 h. Cells were then collected by centrifugation (20 min, 3000g, 4 °C), and the bacterial pellets were frozen at −80 °C until processing. After thawing, bacterial pellets were resuspended in lysis buffer (TrisHCl 50 mM pH 7.5, NaCl 300 mM, glycerol 5%, dithiotreitol 1 mM). Bacteria were lysed by sonication (VibraCell 75115, Bioblock Scientific), and cell debris was removed by centrifugation (15 min, 17,000g, 4 °C). The clarified lysate was filtered at 0.22 µm, and the proteins of interest were purified by a coupled system consisting in affinity chromatography on an Äkta pure Fast Protein Liquid Chromatography system equipped with a Histrap FF crude 1 ml IMAC (immobilized metal affinity chromatography) column (Cytiva). Equilibration and binding and washes were done with 50 mM Tris pH 7.5, 300 mM NaCl, 5% glycerol, 25 mM imidazole, and elution with 50 mM Tris-HCl pH 7.5, 300 mM NaCl, 5% glycerol, and 500 mM imidazole. Eluate was directly loaded on a gel filtration column (GF S200 Superdex) equilibrated in 50 mM Tris-HCl pH 7.5, NaCl 100 mM, glycerol 5%. Eluted fractions were analyzed by Coomassie Blue–stained polyacrylamide gel and concentration of proteins was determined by Nanodrop quantification.

Enzymatic generation Glc-*O*-JA references was conducted through UGT activity as described in Haroth et al 2019 with the following modifications : Ten microgram UGT protein was incubated in 100 µl reaction mixture (50 µM 11OH-JA or 12OH-JA, 50 mM Tris-HCl pH 7.5, 300 µM UDP-glucose, 1 mg/ml BSA) for 1 h at 30 °C. Enzymatic generation of HSO_4_-JA references was obtained through ST2a activity performed as follows : Twenty microgram of ST2a protein was incubated in 100 µL reaction mixture (50 µM 11OH-JA or 12OH-JA, 50 mM Tris-HCl pH 7.5, 5 µM 3’-phosphoadenosine 5’-phosphosulfate (PAPS), 1 mg/ml BSA) for 30 min at 25°C. For both assays, reaction was stopped by adding 50 µL MeOH containing 0.2% acetic acid. After centrifugation, clarified supernatant was analyzed by LC-MS as described above.

### Statistical analysis

Statistical analysis was performed with GraphPad Prism 10 using either one-way ANOVA followed by Tukey post-hoc test with a 95% confidence interval or *t*-test as indicated in Figures.

## Supplemental Figure S1

Impact of impairement of OH-JA biosynthetic pathways on jasmonate profiles upon leaf wounding or fungal infection. Six week-old Arabidopsis plants of WT plants or mutant lines impaired in the deconjugation (*2ah*), or the direct hydroxylation (*3jao*) or both (*5ko*) pathways were either (A) mechanically wounded or (B) infected with *Botrytis cinerea* spores. Leaves were collected after 3.5 h (wounding) or 3 days post-inoculation with fungus. JAs were extracted, analyzed and quantified by LC-MS/MS. Histograms show means ± SEM of 3 biological replicates indicated with circles. Statistical significance was assessed by one-way ANOVA followed by Tukey post-hoc test with a 95% confidence interval. Different letters above boxplots indicate genotypes that are significantly different (P< 0.05). Two independent experiments were performed with similar results.

## Author contributions

Conceptualization, T.H. and L.M.; Methodology; Investigation, T.H., V.M., J.Z., D.D. and D.S.; Analyzing data; T.H., V.M. L.M.; Writing, T.H.; Funding acquisition, T.H. and L.M.

## Acknowledgements

We are grateful to Prof. Minoru Ueda and colleagues (Tohoku University, Japan) for providing the 11OH-JA references and for valuable discussions. Laurence Herrgott is acknowledged for assistance in protein purification, Pauline Delcros for help in mutant generation and enzyme assays, Lisa Marchand, Dennisse Beltran-Valencia and Manasi Nabar for help in mutant plant analysis.

## Notes

### Competing Interest Statement

The authors have declared no competing interest.

