## Supplemental Figure S1 for "Jasmonic Acid Oxidases (JAO) define a new branch in jasmonate metabolism towards 11OH-jasmonic acid and its glucosylated derivative"

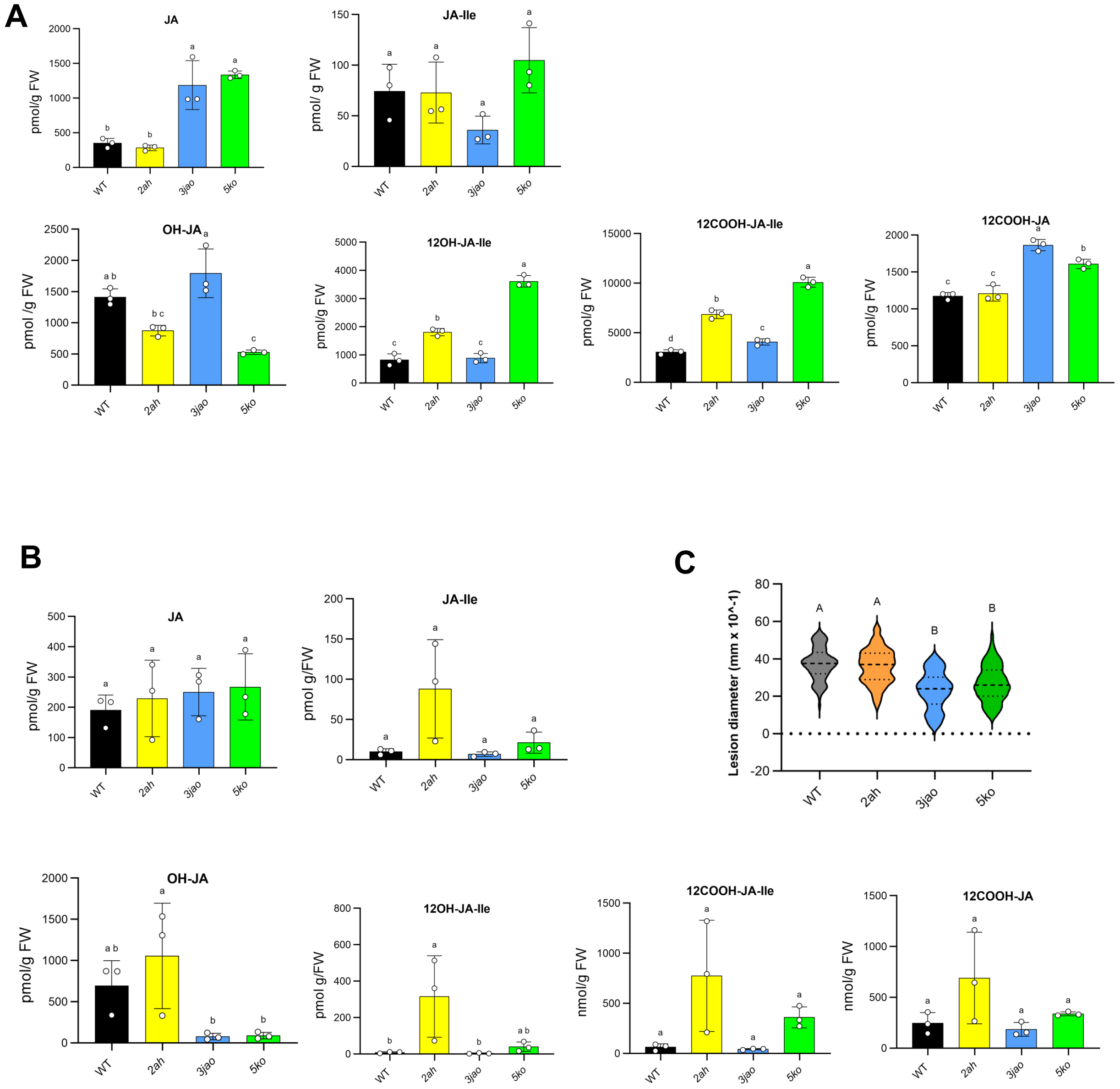


**Supplemental Figure S1**

Impact of impairement of OH-JA biosynthetic pathways on jasmonate profiles upon leaf wounding or fungal infection. Six week-old Arabidopsis plants of WT plants or mutant lines impaired in the deconjugation (*2ah*), or the direct hydroxylation (*3jao*) or both (*5ko*) pathways were either (A) mechanically wounded or (B) infected with *Botrytis cinerea* spores. Leaves were collected after 3.5 h (wounding) or 3 days post-inoculation with fungus. JAs were extracted, analyzed and quantified by LC-MS/MS. Histograms show means ± SEM of 3 biological replicates indicated with circles. Statistical significance was assessed by one-way ANOVA followed by Tukey post-hoc test with a 95% confidence interval. Different letters above boxplots indicate genotypes that are significantly different ($P < 0.05$). Two independent experiments were performed with similar results.
